# APOBEC3A is the predominant global editor of cytosines in human mRNAs and in single-strand RNA viruses

**DOI:** 10.64898/2026.02.10.705151

**Authors:** Zachary W. Kockler, Hamed Bostan, Leszek J. Klimczak, Yun-Chung Hsiao, Matthew S. Dennen, Molly E. Cook, Tony M. Mertz, Ludmila Perelygina, Marat D. Kazanov, Jian-Liang Li, Steven A. Roberts, Dmitry A. Gordenin

## Abstract

APOBEC cytidine deaminases can convert cytosines to uracils in DNA as well as in RNA. The knowledge of DNA deamination motifs preferred by individual APOBECs revealed APOBEC3A as a major source of hypermutation in cancer. However, the extent and relative contribution of specific APOBECs into RNA editing remains unclear as their preferred RNA-editing motifs have not been defined. Here, using a parallel DNA and RNA sequencing strategy, coupled with motif-centered statistical analyses, we sought to identify mRNA edits and diagnostic editing motifs in yeast and human cells overexpressing individual APOBEC enzymes. This approach revealed a prevailing global enrichment for the uCg trinucleotide motif with even greater preference to the motif’s cytosines located in 3’ base of a loop within a hairpin-loop secondary structure when APOBEC3A, but not any other tested APOBEC, was overexpressed. Further analysis revealed the APOBEC3A-like diagnostic motif enrichment in editing calls from human cancers and blood cells. The APOBEC3A-like editing motif also prevailed in the RNA genomes of SARS-CoV-2 pandemic isolates, as well as in infectious persistent rubella viruses, and in polioviruses emerging from live-attenuated vaccine strains. Together, our results indicate that APOBEC3A is the predominant global APOBEC RNA editor with a potential to impact cell physiology and viral evolution.

## Introduction

The apolipoprotein B mRNA editing enzyme, catalytic polypeptide-like 1 (APOBEC1) cytidine deaminase was discovered as a physiologically important programmed site-specific mRNA editor (POWELL *et al*. 1987; TENG *et al*. 1993). Later studies revealed that other related enzymes encoded in the human APOBEC3 gene cluster can also perform untargeted conversion of cytosines into uracils in single-strand RNA as well as in single stranded DNA ((REFSLAND AND HARRIS 2013; LERNER *et al*. 2018; PECORI *et al*. 2022) and references therein). In viral RNA genomes the newly created uracil is maintained and inherited after replication reviewed in (KOCKLER AND GORDENIN 2021). However, for DNA, the results of APOBEC cytidine deamination allow multiple outcomes. If the uracil remains until DNA replication, an adenine base will be placed across it, whereupon further replication will fix the uracil position as a thymine. Additionally, uracil DNA glycosylases can remove uracil bases in DNA to form an abasic site (AP-site), which can ultimately result in any of the 4 nucleotides filling that position (see Figure 1 in (DENNEN *et al*. 2024) and references therein). Altogether, the resulting effects of APOBEC-mediated deamination can lead to DNA hypermutation or RNA hyperediting, which can result in high mutation load in cancers or in antiviral activity (ROBERTS *et al*. 2013; GREEN AND WEITZMAN 2019; ALEXANDROV *et al*. 2020; KOCKLER AND GORDENIN 2021).

**Figure 1.**
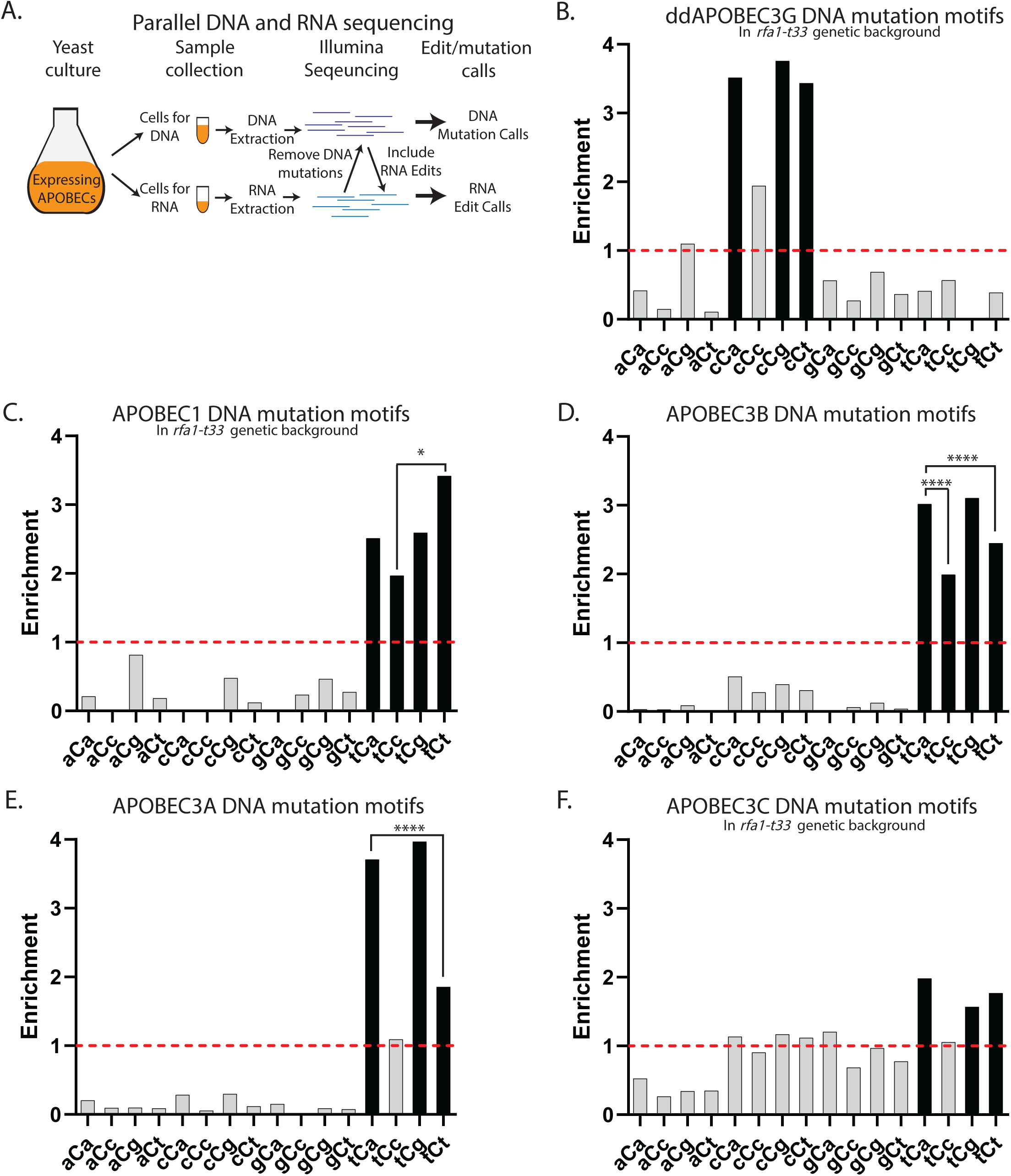
Motif preferred by DNA mutations induced by human APOBECs expressed in yeast. **(A)** Schematic of the parallel DNA and mRNA sequencing strategy for identifying mRNA editing events. Yeast cells are collected from the same culture and processed for DNA and RNA extraction, followed by the whole-genome or whole-transcriptome sequencing . To prevent DNA mutations being called as RNA edits, positions of the DNA genome with at least one read with a mutation were not considered. RNA sequencing reads were aligned to annotated gene transcripts and RNA edits occurring at expected cytosines were identified (see Materials and Methods). (**B-F**) DNA reads from bulk cultures expressing stronger APOBEC3A and APOBEC3B (*RFA1-WT* background) or from *can1* mutants isolated from cultures expressing weaker APOBEC1, APOBEC3C, and ddAPOBEC3G (*rfa1-t33* background) served to call genome-wide mutations, which were used to identify preferred motifs of DNA-mutagenesis by each APOBEC. Enrichments of the 16 possible trinucleotide DNA mutational motifs centered around C to T (including reverse complements) are shown on each panel. Enrichment values shown on each panel can be found in S3 Table. Red dashed lines indicate enrichment level of 1.0. Black bars indicating a statistically significant enrichment (q≤0.05 determined for 16 P-values by Benjamini-Hochberg); grey bars - not significant. Enrichments for all possible trinucleotide mutational motifs, including P-values and q-values can be found in S4 Table. Pairs of odds ratios of all motif-motif combinations within a four-motif uCn group were compared by Breslow-Day Test for homogeneity of the odds ratios (S5 Table). Pairs with statistically significant differences are highlighted by brackets. Level of significance is shown above brackets (*p < 0.05, **p < 0.01, ***p < 0.001,****p < 0.0001). (**B**) ddAPOBEC3G; (**C**) APOBEC1; (**D**) APOBEC3B; (**E**) APOBEC3A; (**F**) APOBEC3C

The roles of different APOBEC family members in DNA mutagenesis was established based on experimentally obtained knowledge of their preferences for oligonucleotide mutational motifs (3-4 nucleotides in size) centered around a mutated cytosine (MERTZ *et al*. 2022). The preferred DNA mutational motif of most of the APOBEC family enzymes is the tCn (where n stands for any base and capitalized C is the deaminated cytosine). The preference to this motif was also clear in two single-base substitution (SBS) mutational signatures (SBS2 and SBS13) agnostically extracted by non-negative matrix factorization (NMF) from large number of mutational catalogs of human tumors (ALEXANDROV *et al*. 2013; ALEXANDROV *et al*. 2020). An exception, APOBEC3G, has a preferred cCn trinucleotide motif (CONTICELLO 2012; REFSLAND AND HARRIS 2013; HARRIS AND DUDLEY 2015; GREEN AND WEITZMAN 2019). The choices of APOBECs potentially responsible for mutating cancer genomes was further narrowed down based upon APOBEC mRNA expression levels in tumor samples where APOBEC3B is the most expressed, while APOBEC3A mRNA correlates best with SBS2/13 loads, suggesting that either or both are responsible for the cancer mutations (BURNS *et al*. 2013a; BURNS *et al*. 2013b). To better determine which APOBEC is responsible for the mutations in cancer, further refinement of the APOBEC mutational motif was performed in yeast ssDNA hypermutation model where either APOBEC3A or APOBEC3B were overexpressed. Preference to mutational motifs were statistically distinguishable in a -2 position upstream the deaminated cytosine. Specifically, APOBEC3A preferred the ytCa motif (Y= pyrimidine) while APOBEC3B had a preference to the rtCa motif (R= Purine). Refined motif preference enabled to classify cancer samples with detectable APOBEC mutagenesis as APOBEC3A-like or APOBEC3B-like. It turned out that the APOBEC3A-like tumors had 10-fold more mutations than the APOBEC3B-like tumors (CHAN *et al*. 2015). Together this suggested that APOBEC3A is the prevailing DNA mutator in human cancers as compared to the other APOBECs. Later, this conclusion was confirmed in direct studies of cultured tumor cells and human tumors (PETLJAK *et al*. 2019; PETLJAK *et al*. 2022; DANANBERG *et al*. 2024; PETLJAK *et al*. 2024). Thus, defining motif preference in a defined yeast model can enable identification of APOBEC enzyme(s) prevailing in cytidine deamination in real life biological contexts.

Initially, APOBEC3 enzymes were considered to be exclusively DNA mutators (REFSLAND AND HARRIS 2013). However, later studies have reported massive editing of cytosines in mRNAs in human cell cultures, as well as in human tumors. Further, editing was increased in response to hypoxia and interferons, in a number of genes important for cell function and for antiviral defense (SHARMA *et al*. 2015; SHARMA *et al*. 2017; SHARMA *et al*. 2019).

Massive RNA editing was detected when APOBEC3A was overexpressed (SHARMA *et al*. 2017) or in tumors with increased APOBEC3A mutagenesis (JALILI *et al*. 2020) indicating capability of this enzyme to serve as a global RNA editor. Another candidate for global mRNA editing was a double-domain APOBEC3G, showing the same as in DNA cCn motif. APOBEC3G-associated edits in cells propagated in atmospheric oxygen concentration were limited to a smaller number of sites; however, edits became more widespread in hypoxic condition (SHARMA *et al*. 2016; SHARMA *et al*. 2019). Both, APOBEC3A and APOBEC3G, showed preference to loops in stem-loop RNA secondary structures and APOBEC3A had even greater preference to cytosines on the 3’-border of the loop (SHARMA AND BAYSAL 2017; JALILI *et al*. 2020). There is a single report on potential mRNA editing by overexpressed APOBEC3B (ALONSO DE LA VEGA *et al*. 2023).

Since this gene is overexpressed in many tumors as well as in normal tissues it should be also included into the list considered as possible global RNA editors in vivo. While there is no data suggesting RNA editing by other APOBEC3 enzymes, their role cannot be excluded *a priori*.

APOBEC enzymes were also implicated into the editing of viral RNA genomes. We found earlier that most of base substitutions in hypermutated vaccine-derived rubella virus isolates from granulomas of individuals with primary immunodeficiency were C to U in plus (genomic) strand. Up to 30% of C to U changes occurred in the uCn trinucleotide motifs conforming with a tCn motif preferred by most of APOBECs in DNA, while the cCn motif preferred by APOBEC3G was depleted (PERELYGINA *et al*. 2019). Further, RNA editing in infectious SARS-CoV-2 single-strand RNA suggested APOBEC’s role prompted by uCn as the primary editing motif for cytosines (DI GIORGIO *et al*. 2020; KLIMCZAK *et al*. 2020; AZGARI *et al*. 2021; RATCLIFF AND SIMMONDS 2021; YANG *et al*. 2025). In model experiments, transient transfection of cultured human cells containing SARS-CoV-2 with an APOBEC3A expression vector caused greater levels of C➔U mutations in uCn motif compared with other APOBEC vectors (NAKATA *et al*. 2023). However, the identity of the specific APOBEC(s) responsible for the bulk of mutagenic editing in genomes of RNA viruses infecting humans have not been revealed.

In both cases, mRNA editing and mutagenesis in the genomes of RNA viruses, more precise knowledge of specific oligonucleotide motifs preferred by different APOBEC enzymes can clarify their relative roles in editing. Here, we utilized a parallel RNA and DNA sequencing approach to reliably identify induced mRNA edits within yeast and human cells that hyperexpressed functional human APOBECs. These mRNA edits were then coupled with knowledge-based motif-centered statistical analyses to quantitatively define the preference of editing motif of each APOBEC. This strategy revealed that though each tested APOBEC can mutate DNA as well as edit RNA, APOBEC3A is the strongest RNA editor. We used the APOBEC3A-like RNA editing motif to assess its prevalence and enrichment within C➔U RNA edits of cellular and viral RNAs. For that purpose, we explored published datasets of mRNA editing in human cancers and blood cells as well as in RNA virus genome editing for vaccine derived rubella viruses (VDRV), polioviruses, and in SARS-CoV-2. We found that in all the biological systems which we analyzed, there was an APOBEC3A-like motif preference, altogether, suggesting that APOBEC3A is not only the prevailing DNA mutator in human cancers, but also predominant global RNA editor among multiple APOBEC enzymes.

## Results

### Experimental design for parallel detection of APOBEC-induced mutagenesis in DNA and mRNA editing in yeast

There are many contributing factors that make identifying mRNA edits more difficult than DNA mutations (reviewed in (KOCKLER AND GORDENIN 2021)). Firstly, each RNA transcript is transient within the cell so a sequencing run will generate a “snapshot” of the current mRNA molecules and any change introduced by editing would disappear when the edited mRNA molecule seizes to exist. On the contrary changes (mutations) in genomic DNA persist and replicate. Secondly, each editing change in mRNA would exist in a small fraction of transcripts within a cell, while genomic DNA mutations are present in 50% or even 100% allelic fraction in a diploid or in a haploid genome, respectively. Moreover, DNA mutations are transmitted to mRNA transcripts and thus obscure RNA edit calling. Altogether, detection of genome-wide (global) RNA editing requires highly accurate sequencing, large sequence coverage, and high efficiency of DNA mutations filtering. To facilitate accurate detection of mRNA edits induced by functional human APOBECs in the budding yeast, *Saccharomyces cerevisiae*, we utilized a strategy of a parallel DNA and RNA sequencing (LERNER *et al*. 2021) (Fig. 1A and Material and Methods). This strategy included overexpressing functional APOBECs in our yeast strains for just 1-2 cell generations followed by parallel DNA and RNA extraction from the same culture (Fig. 1A) and Illumina sequencing at high coverage (target coverage at least 200x for DNA as well as for RNA). For DNA, the high coverage facilitated detection of *de novo* mutations that were present in small fraction of cells, while in RNA it gives the greatest possibility to detect rare editing events. In order to reduce a chance of false calls, high quality RNA edits were called only at positions where cytosines are expected in the RNA transcripts with at least two independent instances of a cytosine to uracil edit. Further, these RNA edits were then cross-referenced with DNA sequencing to remove any position in the transcriptome where there was at least one DNA read with a mutation in the same position as the suggested RNA edit. This limited the possibility of DNA mutations that are transcribed into mRNA and incorrectly called as RNA edits.

The first requirement for the use of an APOBEC ORF for detection of mRNA editing in yeast was a production of deaminase with the *in vivo* enzymatic activity. For that purpose, we relied on detection of DNA mutagenesis caused by expression of that ORF. In our prior study, we assessed the *in vivo* DNA mutagenesis in yeast by every human APOBEC with demonstrated in vitro deaminase activity (DENNEN *et al*. 2024). We found statistically significant mutagenesis in the *CAN1* loss-of-function mutation reporter caused in wildtype yeast by full size APOBEC3A, APOBEC3B ORFs, and by partial single-domain APOBEC3G. We also documented mutagenesis caused by APOBEC1 and APOBEC3C in strains carrying *rfa1-t33* a hypomorph mutation in replication protein A (RPA) subunit which does not bind to DNA as readily as a wild-type RPA. Increased mutagenesis in *rfa1-t33* background was likely due to RPA impeding the access for APOBECs to cytosines in ssDNA ((WONG *et al*. 2021) and references therein). Our prior studies of APOBEC3G mutagenesis in yeast were performed with the partial single-domain APOBEC3G ORF (CHAN *et al*. 2012; CHAN *et al*. 2013), however RNA editing activity in human cells have been reported for full-size double-domain (dd) APOBEC3G (SHARMA *et al*. 2016). Thus, the full-size ORF was used in this study to assess mRNA editing motif preference. Since RPA does not bind efficiently to mRNA as compared to ssDNA (MAZINA *et al*. 2020), we do not anticipate any effect of *RFA1* genotype on RNA editing by APOBECs.

We found that mutagenesis by ddAPOBEC3G increased around 4-fold in *rfa1-t33 vs RFA1-WT* yeast and was also 3-fold increased over empty vector control (S1 Fig., S1 Table). Therefore, parallel detection of DNA mutagenesis and mRNA editing for full size ddAPOBEC3G ORF was also performed in *rfa1-t33* strains. Mutagenesis by APOBEC1, APOBEC3C and ddAPOBEC3G was detectable in *rfa1-t33*, however it was lower than by APOBEC3A or APOBEC3B. Thus, we sequenced DNA from bulk cultures expressing these weaker APOBECs for parallel analysis with RNA reads, but we also sequenced genomes of single colony Can-R isolates for determining spectra of DNA mutations (see Materials and Methods).

### Motif preference by APOBEC-induced DNA mutations in yeast and in human cancers

Using our parallel analysis strategy (Fig. 1A and Materials and Methods), a range of 200-2,000 DNA mutations were called in each group of yeast cultures expressing APOBEC3A, APOBEC3B, ddAPOBEC3G (double-domain), APOBEC1 and APOBEC3C (the latter three in a sensitized *rfa1-t33* background) (S2 Table). These mutations were used to determine motif preferences of DNA mutagenesis caused by human APOBECs in replicating yeast cells. We used knowledge-based motif-centered statistical analyses to calculate and statistically evaluate enrichments all possible dsDNA 96 trinucleotide motifs (including reverse complements) or 192 trinucleotide ssDNA motifs in DNA mutation calls from each data cohort (S3 and S5 Tables).

Since APOBECs cause mutations by C to U deamination in ssDNA as well as in RNA, we then concentrated here and all subsequent analyses of DNA mutagenesis and RNA editing on sixteen trinucleotide motifs centered around C➔T (or C to U) changes in strains expressing each APOBEC (Fig. 1B-F and Materials and Methods). The ddAPOBEC3G that had enriched DNA mutational motifs within the group of four cCn trinucleotides (Fig. 1B), which was consistent with previous studies of APOBEC3G mutagenesis in yeast (CHAN *et al*. 2012; TAYLOR *et al*. 2013). Each of the other four APOBECs tested had enriched trinucleotide DNA mutational motifs only within the group of four tCn trinucleotides (Fig. 1C-F), which is in agreement with prior observations for DNA mutagenesis with these enzymes (summarized in (MERTZ *et al*. 2022)). Enrichment with motifs characteristic for each individual enzyme indicated that each of these proteins, even the weakest APOBEC1 and APOBEC3C had cytidine deaminase activity in cultures where RNA editing activity and motif specificity was aimed to be accessed (Fig. 1A).

There were significant differences for enrichments within four-motif tCn group of APOBECs (Fig. 1C-F). Importantly, APOBEC3A and APOBEC3B preferences of tCa over tCt motif was in agreement with prior studies of APOBEC mutagenesis in yeast subtelomeric ssDNA formed by the muti-kilobase 5’ to 3’ resection as well as in APOBEC mutation clusters in formed in transient stretches of ssDNA in cancer genomes (CHAN *et al*. 2015; SAKOFSKY *et al*. 2019). In the above cited studies APOBEC3A and APOBECB motif-centered analysis was limited to tCa and to the two-motif group tCw (tCa and tCt; w=A or T). We did not include tCc and tCg in prior analyses, where mutagenesis in yeast ssDNA was compared with genome-wide mutagenesis in cancers, because the tCc was likely to overlap with cCn mutagenesis – a motif preferred by APOBEC3G and the tCg with cytosine deamination in meCpG mutagenesis (nCg motif). However, these concerns are not valid for motif analyses in yeast expressing either APOBEC3A or APOBEC3B, because meCpG mutagenesis or APOBEC3G mutagenesis would not be expected. Therefore, we extended analysis of yeast ssDNA mutation catalogs obtained in (CHAN *et al*. 2015) to all sixteen motifs centered around C to T substitution Fig.2 A-C). In that work, we also established preference of APOBEC3A to extended motif ytCa (y=C or T), while APOBEC3B has a preference to rtCa (r=A or G). This enabled to classify APOBEC hypermutated TCGA; ICGC and PCAWG tumors to APOBEC3A-like and APOBEC3B-like (CHAN AND GORDENIN 2015; SAKOFSKY *et al*. 2019; CONSORTIUM 2020). We have also demonstrated that in cancers C- or G-strand coordinated clusters containing more than 3 mutations are highly enriched with APOBEC mutational motifs (Note: based on APOBEC motifs enrichment, G-strand coordinated clusters have originated from cytosine deamination in ssDNA formed by complementary strand). Thus, we used mutation catalogs of C- or G-coordinated long mutation clusters from APOBEC-hypermutated cancer types subdivided to APOBEC3A-like and APOBEC3B-like subgroups of tumors to evaluate enrichments with all sixteen motifs (Fig. 2D,E). In both yeast subtelomeric ssDNA and in long ssDNA hypermutated in cancers, enrichment was observed only in motifs belonging to tCn group and tCa enrichments exceeded enrichments with tCt motif, which was similar to motif-specificity in replicating yeast (compare Fig. 1D,E and Fig.2B-E). However, unlike in replicating yeast, tCa enrichment statistically significantly exceeded tCg enrichment for APOBEC3A-induced ssDNA mutations in yeast and in cancers (Fig. 2B,D) and for APOBEC3B-induced ssDNA mutations in cancer mutation clusters (Fig. 2E). Differences with replicating yeast can be explained by several factors, including greater amounts or strength of RPA binding to short in size and in persistence ssDNA gaps formed during DNA replication as compared to long and persistent ssDNA formed in uncapped telomeres or at yet unknown points of tumor development (see (DENNEN *et al*. 2024) and therein). There was only slight, albeit statistically significant excess of tCg over tCa enrichment for APOBEC3B-induced mutations (2.76116 vs 2.76062, respectively) in yeast subtelomeric ssDNA system (Fig. 2C), which could be due to p-value inflated by high counts of mutation calls and/or genomic background.

**Figure 2.**
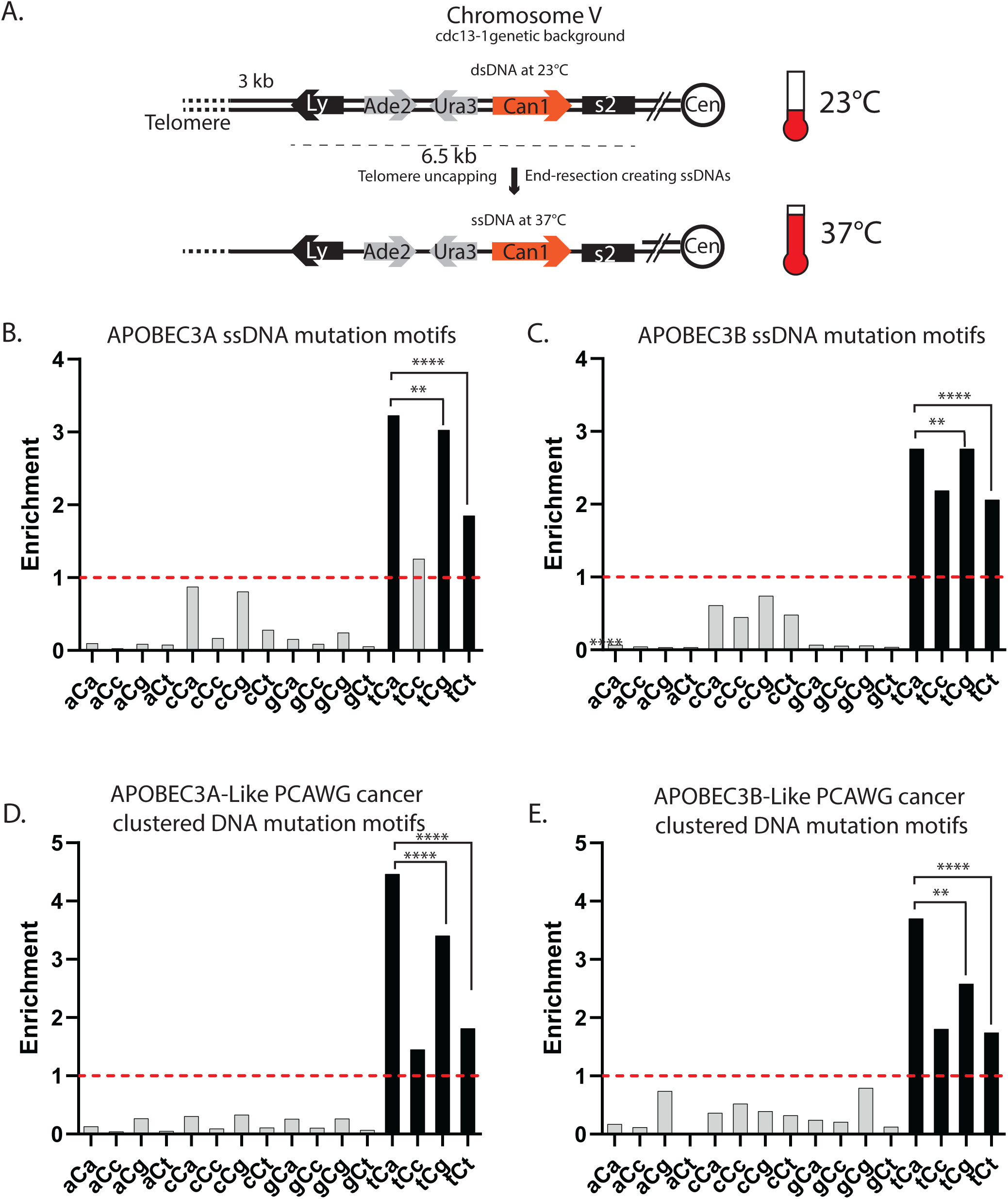
Motif preference for APOBEC3A and APOBEC3B mutagenesis in ssDNA formed in yeast and in human cancers. **(A)** Schematic of the sub-telomeric triple reporter system in (CHAN *et al*. 2015) that exposes long stretches of ssDNA upon the introduction of the 37°C non-permissive temperature. (**B-E**) Enrichment of the 16 possible C to T trinucleotide DNA mutational motifs and statistical evaluations are presented in the same format as in Fig 1B-F. Enrichment values shown on each panel can be found in S3 Table. Red dashed lines indicate enrichment level of 1.0. Black bars indicating a statistically significant enrichment (q≤0.05 determined for 16 P-values by Benjamini-Hochberg); grey bars - not significant. Enrichments for all possible trinucleotide mutational motifs, including P-values and q-values can be found in S6 Table. Pairs of odds ratios of all motif-motif combinations within a four-motif uCn group were compared by Breslow-Day Test for homogeneity of the odds ratios (S5 Table). (**B**) Human APOBEC3A, and (**C**) human APOBEC3B expressed in the sub-telomeric system expressed in the sub-telomeric ssDNA system (data from (CHAN *et al*. 2015)). Note: while tCg enrichment (2.76116) only slightly exceeded tCa enrichment (2.76062), statistically significant p-value (p=0.0048, S5 Table) could be inflated by high counts of observed events. (**D**) Enrichment of the 16 possible C to T DNA mutational motifs in PanCancer Analysis of Whole Genomes (PCAWG) tumors with an APOBEC3A-like whole-genome mutation motif preference and (**E**) with an APOBEC3A-like mutation motif preference (data from (ALEXANDROV *et al*. 2020; CONSORTIUM 2020)).

### Motif preference of APOBEC induced mRNA editing in yeast

We next performed high coverage mRNA sequencing from the same yeast cultures in which APOBEC mutagenesis in DNA was explored (Fig. 1A). Stringent filtering of calls was set to exclude any potential contamination with low-VAF (variant allele frequency) mutation calls in DNA. For that purpose, mRNA C to U change positions in which there was at least a single DNA read containing the matching mutation were excluded. In addition, C to U edits were called as edits only if they were detected in at least 2 reads. We found 8,000 – 10,000 potential mRNA C to U edits in each set of cultures overexpressing APOBEC3A, APOBEC3B, ddAPOBEC3G, APOBEC1 and APOBEC3C (S2 Table and Materials and Methods). We then calculated enrichment for each of sixteen trinucleotide motifs centered around C to U mRNA edits. The C to U mRNA changes in cultures expressing weaker editors - APOBEC1, APOBEC3C, and ddAPOBEC3G - showed small albeit statistically significant enrichments in 6-8 out of 16 motifs (Fig. 3A-C). Only one of these motifs in ddAPOBEC3G expressing yeast fell into the cCn group exclusively enriched in ssDNA of ddAPOBEC3G cultures (compare Fig. 1B and Fig. 3A). The two tCn-specific enzymes, APOBEC1 and APOBEC3C also did not show exclusive enrichment for the group of uCn motifs in RNA (see Fig. 1C vs Fig. 3B and Fig. 1F vs Fig. 3C). Thus, we cannot reliably assign C to U RNA changes to deamination by any of these weaker APOBECs.

**Figure 3.**
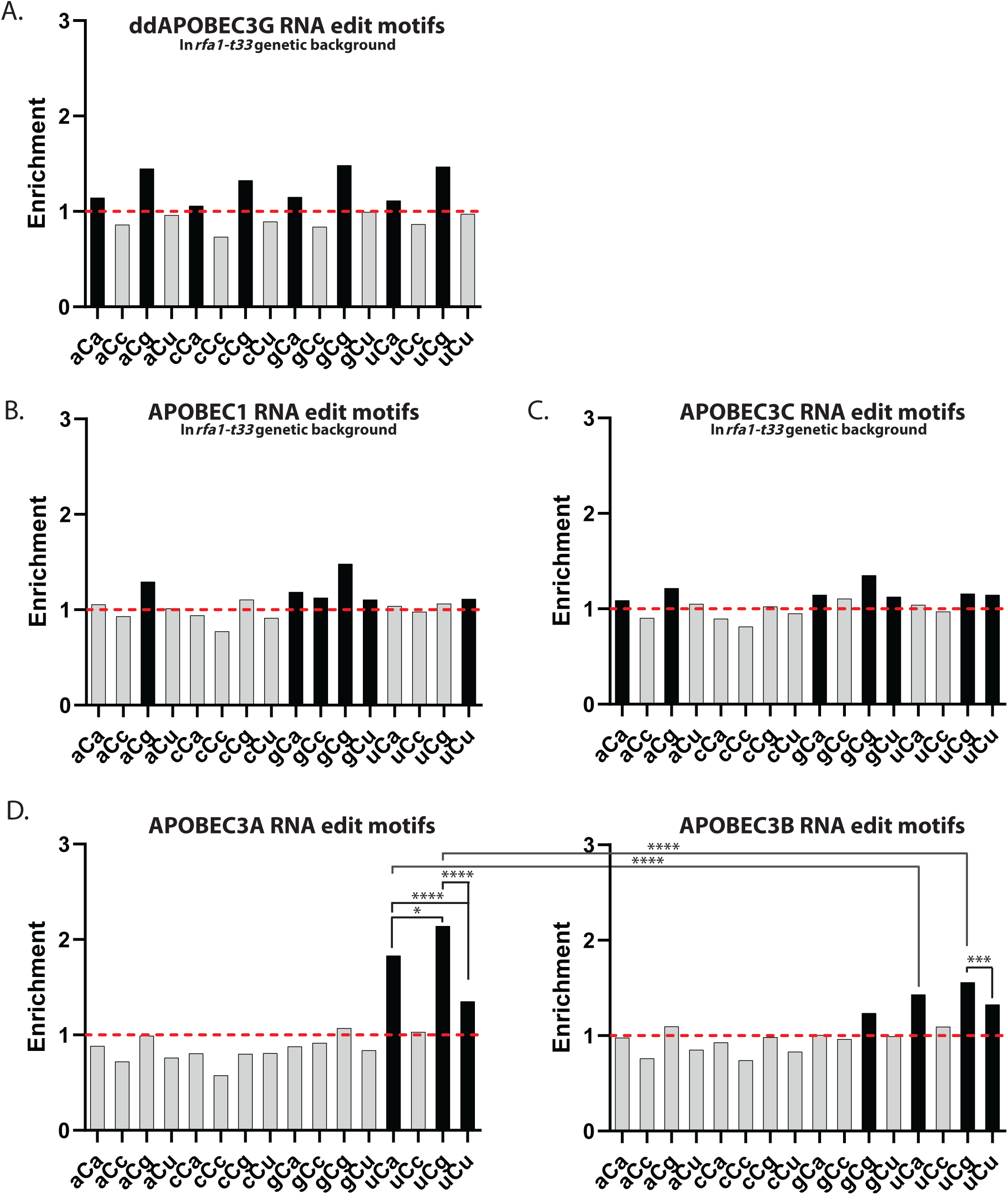
Motif preference of APOBEC-induced mRNA edits in yeast. (**A-D**) Enrichment of the 16 possible C to U trinucleotide mRNA editing motifs and statistical evaluations are presented in the same format as in Fig 1B-F for DNA editing motifs. Enrichment values shown on each panel can be found in S3 Table. Red dashed lines indicate enrichment level of 1.0. Black bars indicating a statistically significant enrichment (q≤0.05 determined for 16 P-values by Benjamini-Hochberg); grey bars - not significant. Enrichments for all possible trinucleotide mRNA editing motifs, including P-values and q-values can be found in S7 Table. Pairs of odds ratios of all motif-motif and cohort-cohort combinations (APOBEC3A vs APOBEC3B) within a four-motif uCn group were compared by Breslow-Day Test for homogeneity of the odds ratios (S5 Table). (A) ddAPOBEC3G. (**B**) APOBEC1. (**C**) APOBEC3C. (**D**) APOBEC3A (left panel) and APOBEC3B (right panel).

Unlike the weaker APOBECs, C to U mRNA changes in yeast expressing APOBEC3A or APOBEC3B showed the same strong (for APOBEC3B) or exclusive (for APOBEC3A) preference to the group of uCn editing motifs selfsame with the tCn group of mutational motifs exclusively enriched in genomic DNAs of strains used for parallel detection of APOBEC DNA mutagenesis and RNA editing. (Compare Fig. 1D,E vs Fig 3D). Thus, RNA-specific base substitution calls in APOBEC3A- and APOBEC3B-expressing yeast can be ascribed to RNA editing with greater confidence. Importantly, there were clear differences between APOBEC3A vs APOBEC3B mRNA editing motif enrichments (Figure 3D). Firstly, mRNA edits in APOBEC3A-expressing yeast showed a clear increase in uCa and uCg enrichments over mRNA edits by APOBEC3B. Secondly, uCg enrichment of RNA edits was statistically significantly greater than uCa enrichment in APOBEC3A but did not show statistically significant excess in APOBEC3B-overexpressing yeast. Importantly, statistically significant excess for preference to uCg over uCa in APOBEC3A yeast mRNA edits (Fig. 3D, left panel) was opposite to statistically significant preference of tCa over tCg in mutagenesis within persistent subtelomeric ssDNA in yeast and in APOBEC3A induced mutation clusters in human cancers (Fig. 2B). The highly significant preference of tCa vs tCg was also found for APOBEC3A-like strand-coordinated DNA mutation clusters in APOBEC-hypermutated human cancers (Fig. 2D).

Establishing the uCg over uCa preference in C to U mRNA changes of APOBEC3A overexpressing yeast and the lack of such a preference in APOBEC3B overexpressing yeast can aid in future analysis of mRNA edits in various biological contexts. Should this difference hold in APOBEC3A and APOBEC3B overexpressing human cells, this could aid in ascribing a specific APOBEC enzyme to RNA editing.

### Motif and structural preferences in mRNA editing by APOBEC enzymes expressed in human cells

After establishing that human APOBEC3A and APOBEC3B not only caused DNA mutations but also can act as mRNA editors with distinguishable motif specificities we explored mRNA editing by these APOBECs in human cells. For that purpose, we expressed human APOBEC3A (A3A-expression) or APOBEC3B (A3B-expression) in human BT-474 breast cancer cell lines and utilized a parallel sequencing strategy similar to the design with yeast cell cultures (Materials and Methods, Figure 1A). Stringent filtering of mRNA changes resulted in sets of 3,000 to 4,000 C to U changes in mRNA of APOBEC-expressing cell lines (S2 Table and Material and Methods). Motif analysis revealed that enriched motifs for both A3A and A3B primarily fell into uCn motifs (with one exception in A3B expressing cell line. There was striking similarity with mRNA editing caused by these enzymes in yeast (compare Fig. 3D and Fig. 4A). Firstly, enrichment with uCn motifs was significantly higher in A3A vs A3B. Secondly, there was strong preference for uCg over uCa mRNA editing in A3A expressing cell line.

**Figure 4.**
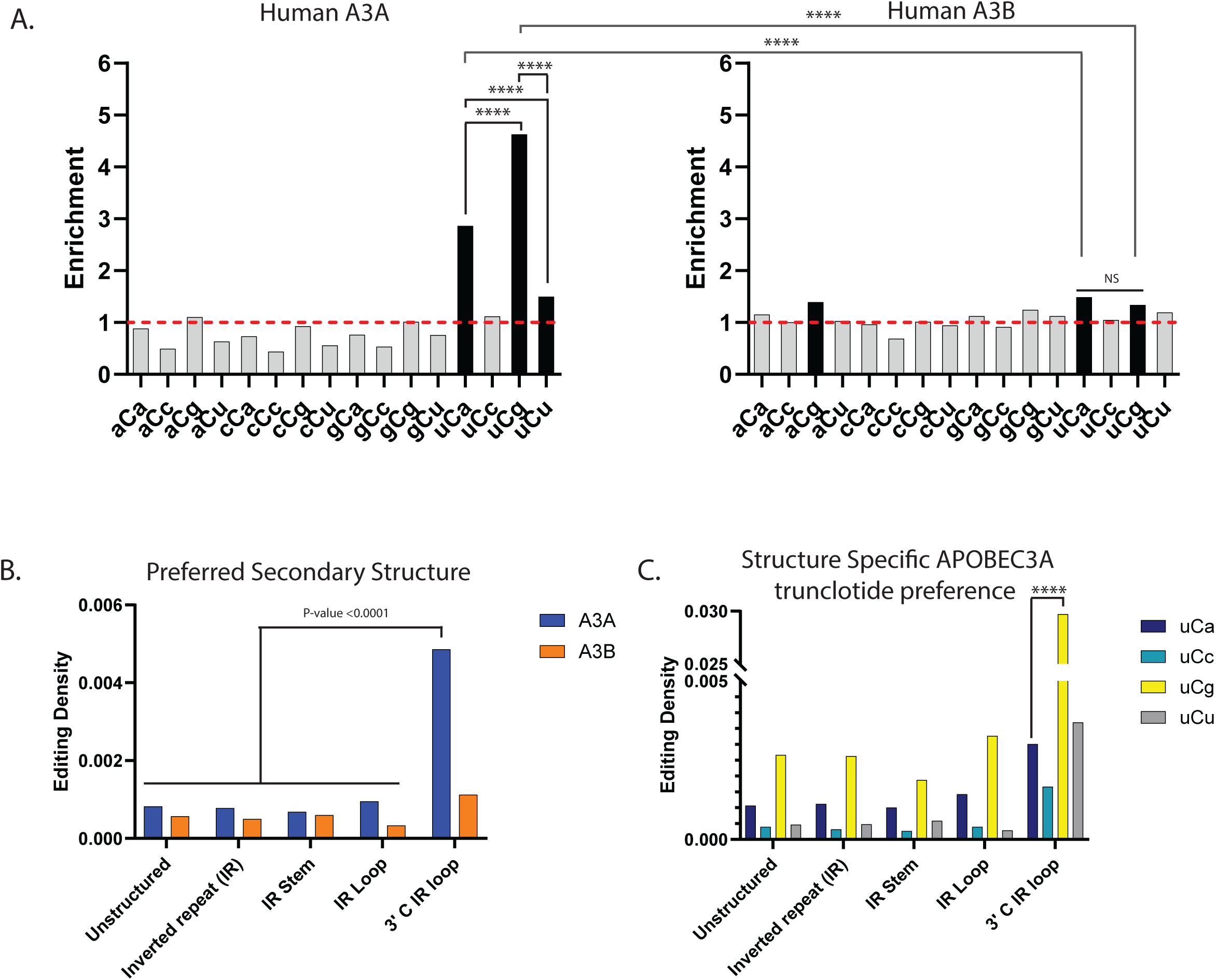
Analysis of APOBEC induced mRNA editing in human breast cancer cell line. **(A)** Enrichment of the 16 possible C to U mRNA editing trinucleotide motifs in human BT-474 breast cancer cell lines expressing APOBEC3A (left panel) or APOBEC3B (right panel). mRNA editing motifs and statistical evaluations are presented in the same format as in Fig 1B-F for DNA editing motifs. Enrichment values shown on each panel can be found in S3 Table. Red dashed lines indicate enrichment level of 1.0. Black bars indicating a statistically significant enrichment (q≤0.05 determined for 16 P-values by Benjamini-Hochberg); grey bars – not significant. Enrichments for all possible trinucleotide mRNA editing motifs, including P-values and q-values can be found in S7 Table. Pairs of odds ratios of all motif-motif and cohort-cohort combinations (APOBEC3A vs APOBEC3B) within a four-motif uCn group were compared by Breslow-Day Test for homogeneity of the odds ratios (S5 Table). (**B**) The density of APOBEC uCn mRNA editing motif in different secondary structure context in cell lines expressing APOBEC3A (A3A), and APOBEC3B (A3B). (**C**) Densities of each of the four APOBEC3A mRNA editing motifs belonging to the four-motif uCn group in each type of secondary structure. (**B,C**) Brackets show significant differences between densities calculated by chi-square test. Source data for panels B and C and statistical comparisons can be found in S8 Table.

It was previously established that human APOBEC3A has preference to 3’C in the loop of stem-loop structures formed by inverted repeats (IRs) in RNA as well as in DNA (BUISSON *et al*. 2019; JALILI *et al*. 2020; OH AND BUISSON 2022). We found similar preference for C to U mRNA edits in A3A but not in A3B cell line (Fig. 4B). Moreover, A3A edits in uCg had highest densities in 3’C positions of IR loop as compared with other uCn motifs (Fig. 4C), which is also consistent with suggestion that most of mRNA edit events in A3A-expressing cell line indeed were generated by A3A.

### C to U transcriptome-wide mRNA editing motif preference in human cancers and in blood cells indicates prevailing role of APOBEC3A

The prevailing role of APOBEC3A in generating genome-wide mutation load in cancer genomes have been established in several studies using the role of germline genotype and distinct motif preference of this enzyme in cytosine deamination in DNA (NIK-ZAINAL *et al*. 2014; CHAN *et al*. 2015; PETLJAK *et al*. 2022). Moreover, the in vitro preference of APOBEC3A to the loop in stem-loop structures is also detectable in APOBEC3A hypermutated cancers (BUISSON *et al*. 2019; PONOMAREV *et al*. 2022). When the preference to loops in RNA stem-loop structures have been explored in whole-exome sequenced tumors from TCGA cohort, such a preference was distinctly associated with APOBEC-hypermutated cancer samples (JALILI *et al*. 2020). We applied motif-centered analysis to over 59,000 transcriptome-wide RNA editing events in a previously reported mixed cohort of 25 tumors with high and 25 tumors with low levels of genome wide APOBEC mutagenesis (JALILI *et al*. 2020). Indeed, enrichment analysis conformed with both features of APOBEC3A RNA editing established in our current study - there was exclusive enrichment in uCn group of motifs and the highest enrichment within this group was with uCg motif (Fig. 5A).

**Figure 5.**
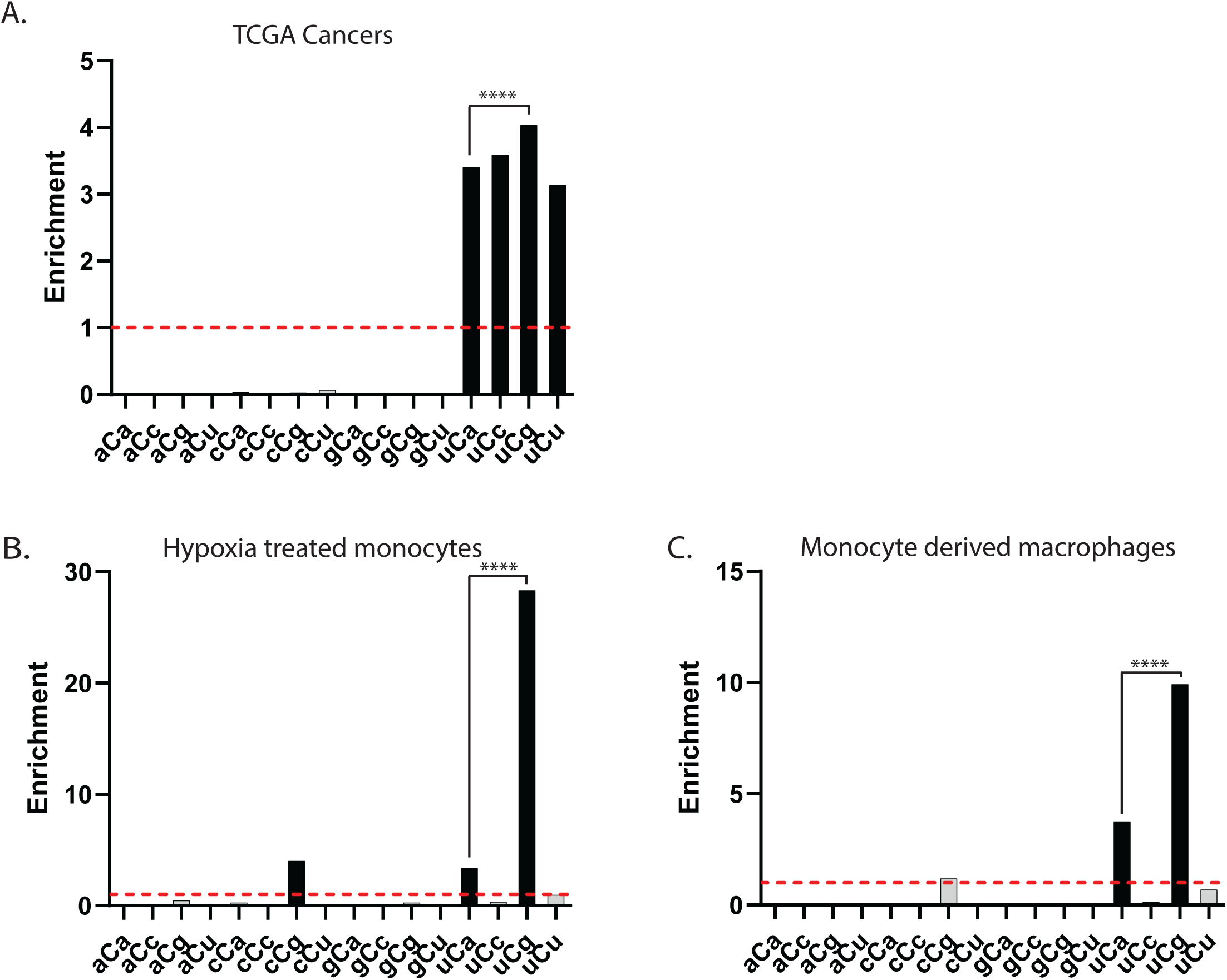
Enrichments for trinucleotide mRNA editing motifs centered around C to U changes in TCGA cancers and in normal blood cells. (**A-C**) Enrichment of the 16 possible C to U trinucleotide mRNA editing motifs and statistical evaluations are presented in the same format as in Fig 1B-F for DNA editing motifs. Enrichment values shown on each panel can be found in S3 Table. Red dashed lines indicate enrichment level of 1.0. Black bars indicating a statistically significant enrichment (q≤0.05 determined for 16 P-values by Benjamini-Hochberg); grey bars - not significant. Enrichments for all possible trinucleotide mRNA editing motifs, including P-values and q-values can be found in S7 Table. Pairs of odds ratios of all motif-motif within a four-motif uCn group were compared by Breslow-Day Test for homogeneity of the odds ratios (S5 Table). (**A**) C to U editing motifs of mRNA from cancer genomes identified in (JALILI *et al*. 2020). (**B,C**) C to U editing motifs of mRNA from hypoxia treated monocytes (**B**); and from monocyte derived macrophages (**C**) (Data from (SHARMA *et al*. 2015)).

We next performed motif centered analysis of the dataset of mRNA edits in non-cancerous blood cells (SHARMA *et al*. 2015). This study demonstrated transcriptome-wide C to U mRNA editing in hypoxia treated monocytes and in monocyte-derived macrophages. Since editing in small number of hotpots was correlated with APOBEC3A expression, authors proposed that this enzyme may also prevail in global mRNA editing. We applied motif-centered analysis to datasets of around 3,000 C to U editing events in monocytes and 140 edits in macrophages (S2 Table). In monocytes as well as in macrophages, uCg motif was enriched way more than any of the other 15 other C to U motifs (Fig. 5B,C). Another enriched motif in both datasets was uCa, also falling into uCn four-motif group characteristic of APOBEC mutagenesis in DNA, which was also detected by our analysis in controlled APOBEC expression in yeast and in human cells (Figs. 3 and 4). C to U edits in monocytes also showed enrichment of cCg motif. Interestingly the same motif was the only detectable outside the uCn group in macrophages. This could be due to either broader motif specificity of APOBEC3A or to the presence of the other low-capacity C to U editor. More studies are needed to confirm that cCg is indeed a new editing motif. Altogether, motif centered analysis of global C to U editing calls in mRNAs of human cancers and in human blood cells revealed prevalence of APOBEC3A diagnostic editing features suggesting that this enzyme is a prevailing RNA editor in humans.

### Enrichment with diagnostic uCg motif indicates a prevailing role for APOBEC3A in editing of (+)ssRNA viral genomes

RNA editing is not limited to mRNA and can occur in other RNA species. In RNA viruses that lack DNA intermediates in their life cycles, changes in viral RNA genomes stemming from various sources, including C to U edits by APOBEC enzymes, may affect viral fitness and/or evolution ((KOCKLER AND GORDENIN 2021) and therein). RNA edits in viral genomes are propagated in progeny viruses and thus can be accurately identified by comparison with the genome of original viral strain providing the original genome sequence is known. The most straightforward detections and assignment of base editing to RNA stand can be done with (+)ssRNA viruses (outlined in Figure 1 of (KOCKLER AND GORDENIN 2021)). However, for most human viruses with either RNA or DNA genomes, identifying the progenitor viral genome by sequencing clinical isolates is challenging due to high genetic variability resulting in heterogeneous populations of viral quasispecies (DOMINGO *et al*. 2012; DOMINGO AND PERALES 2018; DOMINGO AND PERALES 2019).

Nevertheless, there are situations where a progenitor RNA viral genome is known. One such scenario occurs, when a live-attenuated (weakened) vaccine virus persists in the host and undergoes RNA editing enabling within-the-host viral evolution. Vaccine-derived rubella viruses (VDRV) were found in individuals with primary immunodeficiency (PERELYGINA *et al*. 2020).

These patients received the live-attenuated rubella vaccine strain RA27/3 prior to diagnosis of primary immunodeficiency. Over time some individuals developed skin granulomas which contained VDRVs with multiple, up to 300 *de novo* mutations within a ∼10 kb (+)ssRNA genome. These mutations occurred between vaccination and collection of granuloma biospecimens. Motif analyses of VDRV base substitutions revealed that the predominant source of ssRNA genome editing were uCn-specific APOBEC cytidine deaminases (PERELYGINA *et al*. 2019). Here, we performed a motif-centered enrichment analysis with each possible trinucleotide motif centered around 586 C-to-U changes in (+)ssRNA VDRVs reported in that study. We found diagnostic features of APOBEC3A RNA editing, including predominant enrichment with uCn motifs and the highest enrichment in uCg motif (Fig. 6A). Moreover, 290 C-to-U edits in three RNA genomes from VDRVs recovered in another study from idiopathic skin granulomas of clinically immunocompetent individuals (WANAT *et al*. 2022) also exhibited clear diagnostic features of APOBEC3A editing (Fig. 6B). In addition, a modest enrichment in aCg motif was observed in both immunocompromised and immunocompetent individuals (Fig.6A,B). It remains unclear whether this is a minor motif of APOBEC3A-mediated editing, or its enrichment is stemming from another, yet unknown editing mechanism.

**Figure 6.**
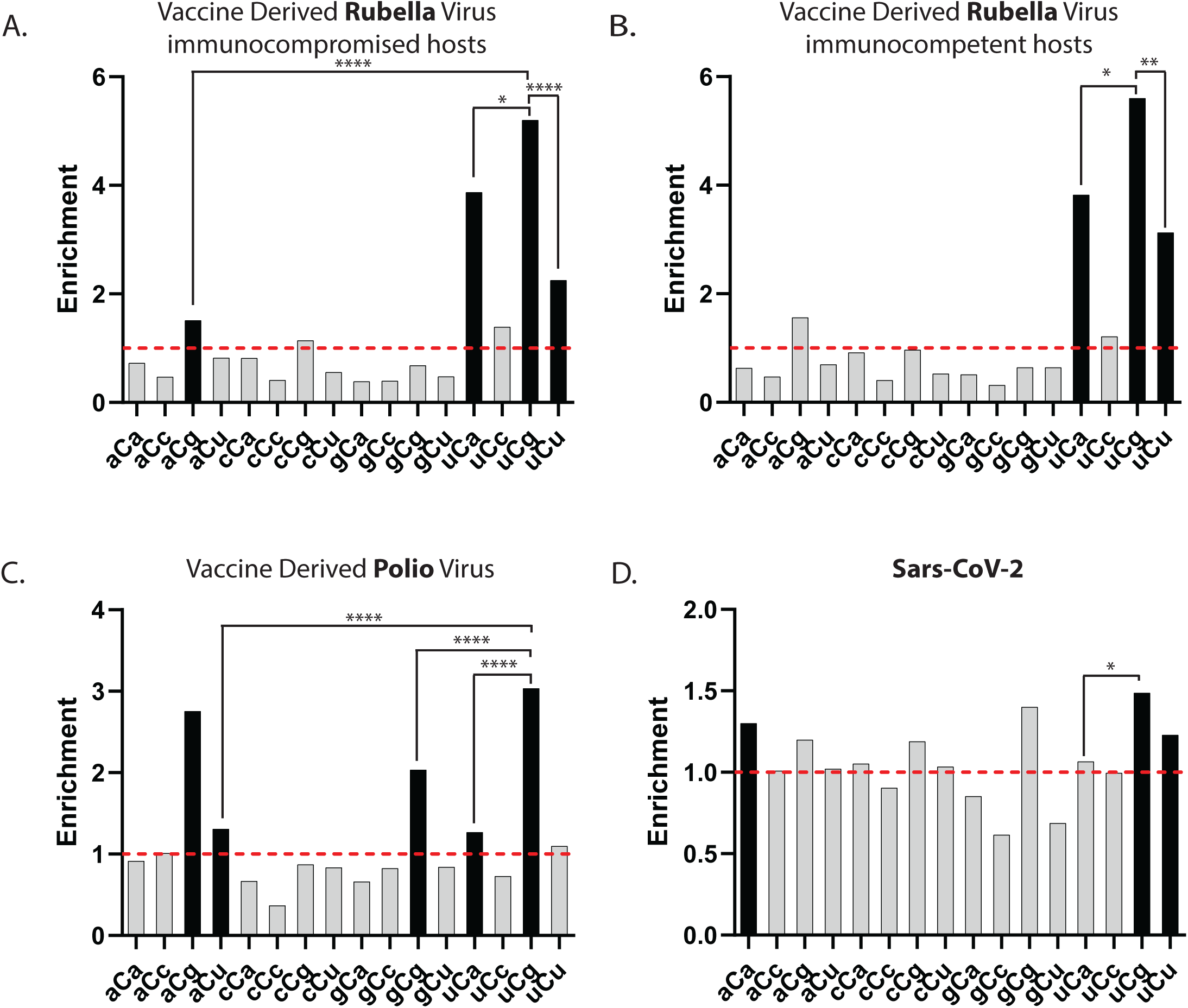
Enrichments for trinucleotide mRNA editing motifs centered around C to U changes in genomes of (+)ssRNA viruses. (**A-D**) Enrichment of the 16 possible C to U trinucleotide mRNA editing motifs and statistical evaluations are presented in the same format as in Fig 1B-F for DNA editing motifs. Enrichment values shown on each panel can be found in S3 Table. Red dashed lines indicate enrichment level of 1.0. Black bars indicating a statistically significant enrichment (q≤0.05 determined for 16 P-values by Benjamini-Hochberg); grey bars - not significant. Enrichments for all possible trinucleotide mRNA editing motifs, including P-values and q-values can be found in S7 Table. Pairs of odds ratios of all motif-motif within a four-motif uCn group and with statistically significant enrichments outside uCn group were compared by Breslow-Day Test for homogeneity of the odds ratios (S5 Table). (**A**) Vaccine-derived rubella from immunocompromised individuals (Data from (PERELYGINA *et al*. 2019)). (**B**) Vaccine-derived rubella from immunocompetent individuals (Data from (WANAT *et al*. 2022)). (**C**) Vaccine-derived polio virus. Accession numbers and publication references can be found in S9 Table. **(D**). Genomes of SARS-CoV-2 pandemic isolates (Data from (KLIMCZAK *et al*. 2020))

The Sabin oral poliovirus vaccine (OPV) is another example of (+)ssRNA live-attenuated virus that can persist and acquire mutations in immunocompromised individuals resulting in emergence of vaccine-derived polioviruses (VDPV). Some nucleotide substitutions in the VDPV genome resulted in reversion to a wild-type phenotype, leading to several poliomyelitis outbreaks in recent years (KEW *et al*. 2004; DEVAUX *et al*. 2023). We analyzed 59 GenBank entries of OPV VDPVs full genome sequences ((CHERKASOVA *et al*. 2002; KEW *et al*. 2002; KEW *et al*. 2004; YANG *et al*. 2005; YAN *et al*. 2010; LIU *et al*. 2015) and unpublished GenBank entries (S9 Table and Material and Methods), which altogether contained 6253 C-to-U edits in (+)ssRNA genomes (Fig. 6C). Similar to rubella VDRV, uCg enrichment exceeded other three motifs of uCn four-motif group. Interestingly, another similarity with rubella virus was statistically significant enrichment with aCg motif not belonging to APOBEC-specific uCn group. Enrichment with aCg was comparable with uCg enrichment (S5 Table; Breslow-Day p-value of difference between two odds ratios = 0.167). Statistically significant enrichments with two other non-uCn motifs, aCu and gCg were much lower than uCg enrichment (Fig. 5C). Altogether, motif-centered analyses of nucleotide substitutions in two vaccine-derived viruses, poliovirus and rubella virus, indicate the prevailing role of APOBEC3A-like editing motif preference with potential involvement of other editor(s).

Another biological context where editing of (+)ssRNA viral genomes in multiple isolates can be traced to a single progenitor strain is represented by sequenced SARS-CoV-2 isolates collected during the COVID-19 pandemic. Here, we analyzed RNA editing events that had the highest likelihood to escape selection pressure, which alter the frequency of variants in pandemic population and thus make the spectrum of editing events recovered from human population to differ significantly from the spectrum of initial editing. For that purpose, we included only C to U edits in non-functional parts of the virus as well as synonymous changes and also counted each individual edit only once. The latter also helped to eliminate confounding effects of edits that have occurred early and propagated in human population though the course of pandemic. Such filtering produced 1375 C to U edits ((KLIMCZAK *et al*. 2020), (S2 Table, Material and Methods, and Fig. 6D). Similar to mRNA edits in model experiments with individual APOBECs-expressing yeast and human cells (Fig. 3D and Fig. 4A), the highest enrichment was observed with uCg motif. There was also statistically significant enrichment with another motif, uCu, belonging to the APOBEC3A-preferred uCn group, as well as with aCa motif, which falls outside this group. Although uCu and aCa enrichments were lower than uCg, there was no statistically significant differences between these two enrichments. Altogether, analysis of SARS-CoV-2 edits is consistent with a prominent role of APOBEC3A in editing (+)ssRNA genomes ; however other editing factors may contribute to aCa motif editing.

In summary, in all biological contexts where C to U RNA editing was identified, including mRNA editing in human cancer, hypoxia treated monocytes, monocyte-derived macrophages mRNA, (+)ssRNA genomes of vaccine-derived viruses, and pandemic population of SARS-CoV-2, we found the best match to the APOBEC3A-like RNA editing motifs previously identified in yeast and human cells that overexpressing APOBEC3A. We therefore conclude that, in addition to being one of the most prolific DNA mutators in cancer, human APOBEC3A is also predominant C to U global RNA editor among multiple APOBEC family enzymes.

## Discussion

In this study, we used a parallel whole genome DNA and mRNA sequencing strategy coupled with a motif-centered statistical analyses to demonstrate that full-size ORFs of APOBEC1, APOBEC3C, APOBEC3G, APOBEC3B, and APOBEC3A expressed in yeast can edit cytosines in mRNA, though to different extents. mRNA editing was evaluated in parallel with DNA mutagenesis. The latter served an internal control of cytidine deaminase activity present in yeast. In order to make DNA mutagenesis detectable even with weaker APOBECs (APOBEC1, APOBEC3C and double-domain APOBEC3G) we used yeast with a hypomorph mutation in the large subunit of RPA (*rfa1-t33*) sensitized to APOBEC mutagenesis (DENNEN *et al*. 2024). The strongest of the editors tested in these defined experiments was APOBEC3A. We also attributed a distinct editing motif of APOBEC3A - uCg→uUg as the most enriched of the trinucleotide motifs centered around C➔U edits. The diagnostic value of uCg RNA editing motif was confirmed in human cells expressing APOBEC3A. The motif preference determined via direct experimental testing and stringent statistical evaluation allowed for the development of a testable hypothesis that APOBEC3A is the primary RNA editor *in vivo*. To test the hypothesis, we applied our knowledge-based statistical analyses to several datasets of RNA editing human mRNAs and in (+)ssRNA viruses and found that the APOBEC3A-like editing motif was prevailing in all the biological datasets that we analyzed. Together, this allowed us to propose that though multiple APOBECs can edit RNA, APOBEC3A is the primary global RNA editor in humans.

Our yeast study revealed several features of C➔U RNA editing that can be attributed to APOBEC3A. (i) The highest total enrichment in the uCn group of four trinucleotides; (ii) lack of enrichment of the remaining twelve trinucleotide motifs centered around C; (iii) unique preference to uCg→uUg editing motif over uCa→uUa (Figure 3). The uCg over uCa preference is unique to RNA editing and was not observed in APOBEC3A mutagenesis in the analogous ssDNA motifs tCg◊tTg and tCa◊tTa in yeast (Figs. 1 and 2). It remains to establish mechanisms underlying motif preferences to APOBEC3A mRNA editing, however the combination of the three diagnostic features held in human cells overexpressing APOBEC3A and was not observed in cells overexpressing APOBEC3B (Figure 4 A). In agreement with prior biochemical studies of APOBEC3A and APOBEC3B (SHARMA AND BAYSAL 2017; CORTEZ *et al*. 2019; JALILI *et al*. 2020), the APOBEC3A expressing human cells demonstrated the strongest preference to the loop of stem-loop structures (Figure 4B) and furthermore, showed clear preference for the 3’ base of a hairpin’s loop in APOBEC3A expressing cells. Moreover, the preference of uCg motif to 3’C in the loop was the highest among four motifs of the uCn group (Figure 4C).

The APOBEC3A-like editing features prevailed in mRNA editing calls from TCGA cancers and human blood cells, where the agent of C➔U global RNA editing was not defined (Figure 5). General preference to uCn motif editing in yeast was observed for APOBEC3B as well as for APOBEC3A, however APOBEC3A-like uCg motif prevailed in cancers and in blood cells. It is surprising that APOBEC3A-like editing appears to prevail over APOBEC3B, because the latter exhibits stronger expression across human cells (BURNS *et al*. 2013a; BURNS *et al*. 2013b). It is worth noting, however, that biochemical studies (XIAO *et al*. 2017; CORTEZ *et al*. 2019) have shown that DNA editing by APOBEC3B was inhibited by RNA, whereas APOBEC3A-mediated DNA editing was not affected.

Multiple variants of mRNA transcripts derived from a single gene can be generated by end processing, non-coding base modifications as well as by alternative splicing and editing events which result in downstream modifications of protein sequence (SHARMA *et al*. 2025).

While site-specific RNA editing and alternative splicing provide additional opportunities for programmed cell differentiation, the dispersed global editing events involving mRNAs and non-coding RNAs may also carry physiological functions (LERNER *et al*. 2018; CHRISTOFI AND ZARAVINOS 2019; GASSNER *et al*. 2020; ALQASSIM *et al*. 2021; DONATO *et al*. 2022; PECORI *et al*. 2022). Background of global, dispersed across multiple sites, mRNA editing would also generate a background of disordered variants of proteins with yet unknown physiological impact at the cell and organism levels (UVERSKY 2002; HOLEHOUSE AND KRAGELUND 2024; UVERSKY 2025). Biological function assigned to global DNA and RNA editing by APOBEC enzymes is the antiviral innate immunity regulated within the large network of innate immunity interferon stimulated genes (ISGs) (HARRIS AND DUDLEY 2015; BABADEI *et al*. 2024), which respond to a variety of environmental and endogenous stimuli. Thus, global mRNA editing may be an inevitable consequence of the existence of complex innate immunity system. Knowledge about relative contributions of individual APOBECs should be included into design of future research of mechanisms generating epitranscriptome. The high input into mRNA editing capability of APOBEC3A revealed by our study suggests a testable hypothesis that can be addressed in human population. It was established that APOBEC3A-APOBEC3B deletion-fusion variant frequent in human population increases frequency of APOBEC3A-like mutagenesis (NIK-ZAINAL *et al*. 2014; CHAN *et al*. 2015; CHEN *et al*. 2019). The facilitated mutagenesis was explained by increased stability of a fusion transcript carrying complete APOBEC3A ORF with 3’-UTR of APOBEC3B mRNA (CAVAL *et al*. 2014). Based on our current study, we anticipate that the overall level of global mRNA editing as well as the enrichment with uCg mRNA editing motif in individuals carrying APOBEC3A-APOBEC3B fusion variants would be greater as compared to wild-type carriers.

Accurate detection of APOBEC-catalyzed mRNA editing by whole transcriptome sequencing is challenging as compared to APOBEC induced mutations in genomes of RNA viruses, because each mRNA editing event should be detected in the same molecule in which the C➔U change occurred, while each mutation in viral RNA genomes can be reliably identified after propagation of mutants. Since structural constraints define exquisite specificity of APOBECs to single-strand polynucleotides, mutagenesis frequency and strand-specificity of APOBEC mutagenesis in RNA viruses would be defined by their replication cycle. The highest frequency of mutant genomes would be expected in (+) ssRNA viruses with C➔U generated in genomic (+) strand (see Figure 1 in (KOCKLER AND GORDENIN 2021)). Viruses with positive ssRNA genomes comprise large fraction of RNA viruses, which are distributed across multiple host species (WARD 1993) including humans carrying single or multiple APOBEC genes (CONTICELLO *et al*. 2005). Indeed, the striking level of APOBEC-induced hypermutation was detected in rubella virus recovered from granulomas of patients with Primary Immunodeficiency (PID) (PERELYGINA *et al*. 2019) and from immunocompetent patients (WANAT *et al*. 2022).

Sequence comparison clearly pointed to the attenuated rubella vaccine virus being a target of hypermutation. This allowed to identify position and surrounding nucleotide context of each individual C➔U mutation event and calculate enrichment with each mutation motif supported by rigorous statistical evaluation (Fig. 6A,B). Two other real-life events in which complete sequence of originating genome was known, vaccine-derived polio virus (LIU *et al*. 2015; DEVAUX *et al*. 2023) and pandemic isolates of SARS-CoV-2 (KLIMCZAK *et al*. 2020; RATCLIFF AND SIMMONDS 2021) enabled rigorous motif enrichment calculation (Fig. 6C,D). In all cases, enrichment with APOBEC3A-like uCg motif was the strongest. This result points to APOBEC3A as the strongest candidate for the global RNA editing not only in mRNAs, but also in viruses with (+)ssRNA genomes propagating in humans. Since APOBEC3A as well as other APOBECs are under innate immunity and inflammation control, it would be important to explore the effect of the innate immunity and inflammation triggers on sequence variation of (+)ssRNA virus(es) during virus persistence or propagation in humans. In support of this suggestion, C➔U editing of SARS-CoV-2 RNA genomes in human cultured cells was indeed facilitated by interferons and by inflammation-related TNF-alfa (NAKATA *et al*. 2023).

While our study presented evidence of prevailing role of APOBEC3A in global editing of cytosines mRNAs and in RNA viruses, this statement is based on a limited amount of data analyses. More studies are required to evaluate the level of generality of such a prevailing role. Regardless, the mRNA editing the APOBEC3A-like uCg→uUg editing motif identified in our work suggests testable hypotheses about impacts of individual genotype and environmental factors on generation of RNA sequence variants in cell models and even in mRNAs of cells from individuals or in RNA viruses in humans.

## Materials and Methods

### Strains and plasmids

The yeast strains were isogenic to ySR128 (ROBERTS *et al*. 2012). Strains were MATalfa mating type carried *his7-2 leu2-3,112 trp1-289* mutations: *CAN1, ADE2, URA3* were deleted from their original positions. In this background the *lys2::ADE2-URA3-CAN1* array was inserted at the native *LYS2* on the right arm of chromosome II, approximately 230 kb from the centromere and 340 kb from the telomere. An additional replacement deletion *ung1::NAT* eliminated uracil DNA glycosylase, thereby preventing creation of AP-sites at positions of cytosine deamination in DNA. One additional strain of the same lineage was created by introducing the RPA DNA binding deficient *rfa1-t33* mutation, which was confirmed by Sanger sequencing. These two strains were transformed with a Hyg^R^ marked autonomously replicating vector with a doxycycline inducible full-size APOBEC inserted ORFs into the empty vector (EV) backbone: pSR4199 (EV), pSR433 (APOBEC1), pZAK031 (APOBEC3A), pSR440 (APOBEC3B), pSR469 (APOBEC3C), pZAK045 (full-size double-domain, ddAPOBEC3G) (MERTZ *et al*. 2025). Yeast strains and plasmids are listed in S10 Table. Autonomous replication status of vectors in transformants was confirmed by frequent loss of Hyg^R^ plasmid marker after propagation on the media without antibiotic. All *RFA1* wild type yeast strains for experiments described in this paper were grown at 30°C and *rfa1-t33* strains were propagated at 23°C except during short-term induction of APOBEC ORFs performed at 30°C.

### Parallel DNA and RNA sample collection in yeast

Maximizing the power of our parallel DNA and RNA sequencing approach required the collection of two yeast samples taken from the same culture in parallel. The yeast strains overexpressing APOBECs were streaked for single colonies on Hyg media directly from -80°C stocks and grown for 2 days at 30°C (*RFA1* wild type yeast) or for 3 days at 23°C (*rfa1*-t33 yeast). The entire single colony (∼1x10e+07 cells) was suspended in 50mL of liquid YPDA media supplemented with Hygromycin 0.2 mg/mL to a density of 2x10e+05 cell/mL and grown at 30°C or at 23°C (*RFA1-WT* or *rfa1-t33*, respectively) with agitation at 250 rpm to a cell density of ∼1.0x10e+07 cells/mL (approximately 12-14 hours). For each APOBEC as well as for empty vector control, 16 to 24 cultures were grown. Once cultures reached designated density, doxycycline was added to final concentration of 10 µg/mL to stimulate APOBEC genes expression. Cells of all strains were then grown at 30°C for 4 hours (1-2 cell division(s)) to stimulate expression of the APOBECs. Then, as quickly as possible, 2mL of cells were aliquoted for RNA extraction while 10mL were aliquoted for DNA extraction. These aliquots were spun down and the liquid was removed by aspiration leaving only the yeast cell pellet. The pellet was then flash frozen in liquid nitrogen and stored in -80°C until extraction (typically within a week). The RNA extraction was performed using the Direct-zol™ RNA MiniPrep from Zymo Research, Irvine CA, USA (Cat # R2052). DNA extraction as performed using the YeaStar Genomic DNA Kit from Zymo Research (Cat # D2002) following the standard chloroform version of the protocol, with the modification of doubling the digestion buffer and lysis buffers volumes. In addition to DNA extraction from bulk cultures, genomic DNA was also extracted from single colonies of canavanine resistant mutants (16 for EV, 24 for APOBEC3C and 36 for ddAPOBEC3G).

### Parallel DNA and RNA sample collection in human cells

#### Cell culture and creation of stable cell populations

BT474 cells were obtained directly from ATCC as part of a breast cancer cell line panel, ATCC 30-4500K. BT474 and derivative cell populations were propagated according to ATCC guidelines in Hybri-Care media with 1.5 g NaHCO_3_ per liter and 10% FBS. Stably transduced BT474 cells were selected with 20 µg/mL blasticidin, 500 µg/mL hygromycin B, or 1 µg/mL puromycin. HEK293T and HEK293T-TETR cells (described in (CORTEZ *et al*. 2019)) were grown in DMEM+10% FBS or DMEM+10% FBS with 5 µg/mL blasticidin, respectively. List of human cells lines can be found in S10 Table.

BT474 derived cell populations were established by serial lentiviral transductions as follows. Lentivirus to establish TETR expression was produced by co-transfection of pLenti_CMV_TetR_Blast (Addgene, Watertown, MA, USA Cat #17492) into HEK293T cells with psPAX2 (Addgene, Cat#12260) and pMD2.G (Addgene, Cat #12259). Lentiviruses for pTM406 (empty vector), pTM664 (APOBEC3A overexpression), and pTM666 (APOBEC3B overexpression) were produced similarly but used HEK293T-TETR cells, in which APOBEC expression is repressed by TETR, which allows more efficient lentivirus production by limiting APOBEC-mediated restriction. To generate BT474 cells with doxycycline inducible empty vector control, APOBEC3A or APOBEC3B, cells were first transduced with pLenti_CMV_TetR_Blast to establish TETR expression. Post selection, these cells were subsequently transduced with either pTM406, pTM664, or pTM666 to introduce the empty vector control, APOBEC3A, or APOBEC3B coding sequences. For transductions, lentiviral containing supernatant was added to BT474 cells in the presence of Lentiblast Premium (OZ Biosciences, San Diego, CA, USA), which was diluted 1:200 in the cell culture media. Following transductions, the media was changed at 16 hours, and selection started at 72 hours. The plasmids pTM664 and pTM666 were previously described (reference, (CORTEZ *et al*. 2019)). pTM406 was constructed from an LR Clonase (ThermoFisher, Waltham, MA, USA Cat# 11791020) reaction between pLenti PGK Puro DEST (Addgene, Cat #19068) and pENTR1A no ccDB (w48-1) (Addgene, Cat #17398) and sequence validated by Sanger sequencing.

#### Human cell lines preparation for next-generation sequencing

Expression of APOBEC3A and APOBEC3B in BT474+pLenti_CMV_TetR_Blast, pTM664 and BT474+pLenti_CMV_TetR_Blast, pTM666 cells were induced by addition of 1.0 µg/mL doxycycline to the cell culture media 96 hours before harvesting cells.

BT474+pLenti_CMV_TetR_Blast, pTM406 cells were similarly treated and used as empty vector controls. RNA and DNA were harvested from asynchronous cell populations at approximately 60% confluence from two 10 cm dishes for each prep. RNA was purified using the E.N.Z.A. Total RNA Kit (Omega Bio-Tek, Norcross, GA, USA, Cat# R6834-00) followed by DNase I treatment, phenol chloroform extraction, and ethanol precipitation. DNA was purified using the DNA Blood and Tissue kit (QIAGEN, Germantown MD, USA). Isolated DNA was subsequently purified from a contaminating nuclease by phenol chloroform extraction and ethanol precipitation. The integrity and purity of isolated RNA and DNA was confirmed by visualization on an agarose gel with 1% vol/vol bleach or standard agarose gel stained with GelRed, respectively.

### DNA mutation calling in yeast

DNA mutations were called after sequencing at the NIEHS epigenomics core facility where DNA libraries were prepared using the Illumina DNA Prep kit and sequenced on the NextSeq or NovaSeq platforms. High mutagenic activity of APOBEC3A and APOBEC3B in yeast even without induction of Tet-promoter allowed to detect sufficient number of APOBEC-induced mutations from whole-genome sequencing of DNA prepared from bulk cultures. Mutagenesis with APOBEC1, APOBEC3C and ddAPOBEC3G was weaker ((DENNEN *et al*. 2024) and S1 Figure), therefore we used sensitized background *rfa1-t33* and prepared DNA from single-cell canavanine-resistant colonies isolated from the same bulk cultures that were used for high-coverage DNA and mRNA sequencing. The Illumina sequencing read files were imported into CLC Genomics Workbench (Version 20.0, QIAGEN, Germantown MD, USA) for read mapping to the ySR128 yeast reference genome (GenBank accessions CP036470.1 - CP036486.1 and (ROBERTS *et al*. 2012)) with duplicate reads being removed. Mutation calls for DNA motif analyses from whole-genome sequencing were obtained using CLC Genomics Workbench. For bulk cultures expressing highly mutagenic APOBEC3A and APOBEC3B calls were obtained using basic Variant Caller option with subsequent filtering to a minimum allelic frequency of 20% in order to limit DNA mutation motif analysis to APOBEC induced mutations occurring during the first1-2 cell divisions of the cell at the origin of the culture. Multistep workflow is detailed at S11 Table. Mutation calls from single-cell canavanine resistant colonies from APOBEC1, APOBEC3C and ddAPOBEC3G culture were obtained using multistep workflow centered around the Fixed Ploidy Variant Detection tool (ploidy of 1) with subsequent filtering to a minimum allelic frequency of 80%. The multistep workflow can be found in S12 Table. For both workflows, the resulting mutations were then filtered to remove calls that occurred in multiple samples to limit analysis independent events. These filtered mutation call datasets were then listed in Mutation Annotation Format (MAF) General description of MAF format is in (https://docs.gdc.cancer.gov/Data/File_Formats/MAF_Format). MAF files were then processed using the P-MACD pipeline (https://github.com/NIEHS/P-MACD) All MAF files fed into P-MACD and their derivative versions *.anz4, containing +/-20 nucleotide flanks around each base substitution are listed in S2 Table and can be found at https://doi.org/10.5281/zenodo.18079216.

### Parallel DNA and RNA sequence reads processing for RNA edit calling

Reads from paired DNA- and RNA-Seq samples (Submitted to SRA: Bio-project PRJNA1398811 for human samples; Bio-project PRJNA1398896 for yeast samples) were evaluated for quality and subjected to trimming to remove adapters and low-quality base sequences using TrimGalore (KRUEGER 2023). High-quality cleaned DNA-Seq reads were aligned to their respective reference genomes (ySR128 (ROBERTS *et al*. 2012) for yeast samples and Human hg19 for human samples) using minimap2 (LI 2018) (version 2.24-r1122) with the parameters --MD -ax sr. Similarly, high-quality cleaned RNA-Seq reads were aligned to their corresponding reference genomes using HISAT2 (ZHANG *et al*. 2021) (version 2.2.1) with the parameters --dta.

Aligned reads from both DNA-Seq and RNA-Seq data (in SAM format) were processed using the Picard toolkit (WAY 2022) (version 2.26.11). The following steps were performed:

1. **AddOrReplaceReadGroups**: Appropriate read groups were added, and SAM files were converted to sorted BAM files with the parameters: VALIDATION_STRINGENCY=SILENT, SO=coordinate, and RGPL=illumina.
2. **MarkDuplicates**: PCR duplicates were marked using the parameters: CREATE_INDEX=true and VALIDATION_STRINGENCY=SILENT.

The resulting paired samples (aligned, sorted, indexed, and duplicate-marked DNA and RNA BAM files) were subsequently used as input for RNA editing analysis with the REDITools (PICARDI 2024) pipeline.

### RNA Editing Calling

RNA edits for both yeast and human were identified using REDItoolDnaRna.py from the REDITools (PICARDI 2024) (version V1) pipeline with the parameters: -s 2 -t 20 -n 0.05 -N 0.05 - e -E -d -D. For human genome hg19 reference was used. The reference annotation ySR128_v2.gff file (https://doi.org/10.5281/zenodo.18079122) for yeast ySR128 genome was used for accurate transcript mapping. DNA positions used to identify RNA editing calls were filtered to positions where only cytosines were present in DNA reads to limit potential DNA mutations being called as RNA edits. RNA edits were only called at positions containing only cytosines in DNA reads while cytosines or thymines were called in those positions. This resulted in the production of the final high-confidence C to U RNA editing call set.

### MAF Files Generation for RNA edits

The final filtered sets of RNA editing calls for yeast and human experiments for each sample were converted into the MAF format for the downstream analysis similar as outlined for DNA mutation calls with additional information relevant for RNA edit filtering included (S2 Table). The RNA editing calls in these MAF files were then filtered to include only editing calls where more than one read had evidence of a cytosine to uracil (displayed as thymine) RNA edit (abbreviated to rTg1). Yeast calls were further filtered to remove empty vector calls from calls in APOBEC overexpressing cells to account for potential calls coming from a non-APOBEC cytosine-deamination source. Human cell experiments were analyzed as without further filtering because empty vector cells contain all APOBECs as well as non-APOBEC sources of potential deamination. Even so, empty vector cells had a low enrichment in motifs characteristic of APOBEC3A or APOBEC3B (see Results). MAF files for mRNA editing events from publications (SHARMA *et al*. 2015; PERELYGINA *et al*. 2019; JALILI *et al*. 2020; KLIMCZAK *et al*. 2020) were generated directly from reported edit calls. MAF files for RNA edits in vaccine-derived isolates of ss(+)RNA viruses, Rubella (WANAT *et al*. 2022) or poliovirus ((CHERKASOVA *et al*. 2002; KEW *et al*. 2002; KEW *et al*. 2004; YANG *et al*. 2005; YAN *et al*. 2010) and unpublished GenBank entries (S9 Table) were generated by comparing GenBank genome sequences of a virus isolate with a GenBank reference of the parent virus, using the procedure described in (PERELYGINA *et al*. 2019; KLIMCZAK *et al*. 2020)

### Datasets of APOBEC-induced mutations in ssDNA

#### APOBEC-induced mutations formed in yeast subtelomeric ssDNA (CHAN et al. 2015)

Weutilized a dataset of mutations induced by either APOBEC3A or APOBEC3B expressed in long stretches of subtelomeric ssDNA formed by 5’ to 3’ resection of *ung1*Δ yeast carrying *cdc13-1* defect in telomere capping (Fig. 2A). Mutation catalogs were taken directly from (CHAN *et al*. 2015) and reorganized as *anz4 version of MAF format (S2 Table). Similar to (CHAN *et al*. 2015; SAINI *et al*. 2020; HUDSON *et al*. 2023) APOBEC-induced mutations in C:G pairs formed at the distance less than 30 kb from the left telomeres were assigned to cytosines of the ssDNA in the unresected bottom strand, while mutations at the distance less than 30 kb from the right telomeres were assigned to cytosines of the ssDNA in the unresected top strand. MAF files with strand assigned APOBEC-induced mutations were then used to calculate enrichments with all possible 192 mutational motifs as described in (KLIMCZAK *et al*. 2020; HUDSON *et al*. 2023).

#### C- of G-strand-coordinated mutation clusters formed in long persistent stretches of ssDNA in human cancers

We utilized the output of APOBEC mutagenesis analysis from Data Resources of Pan-cancer Analysis of Whole Genomes (PCAWG) (see Supplementary Table 4 in *<u>(Consortium 2020)</u>*. Previously, we have demonstrated that several types of human cancers included in PCAWG dataset – Bladder (Bladder-TCC), Breast (Breast-AdenoCA and Breast-LobularCA), Cervical (Cervix-SCC), Head and Neck (Head-SCC), Lung (Lung-AdenoCA and Lung-SCC) (PCAWG abbreviated names shown in parentheses) are highly enriched in APOBEC mutational motif and can be even separated into APOBEC3A-like and APOBEC3B- like tumors by detailed motif analysis (ROBERTS *et al*. 2013; CHAN *et al*. 2015). Moreover, we reveled that in APOBEC hypermutated tumors C- or G-strand-coordinated clusters containing more than three mutations always demonstrate higher enrichment of APOBEC mutational motif than the scattered mutations in C:G pairs. Presence of all other C:G preferring motifs, e.g. the nCg to nTg motif of meCpG mutagenesis were depleted in C- or G-strand-coordinated clusters (ROBERTS *et al*. 2013; ROBERTS AND GORDENIN 2014; CHAN AND GORDENIN 2015; KAZANOV *et al*. 2015; GERHAUSER *et al*. 2018). Therefore, we filtered mutations in APOBEC-hypermutated types of PCAWG cancers to only C:G pair mutations in C- or G-strand-coordinated clusters containing more than three mutations of tumors that can be also assigned to either APOBEC3A- like or to APOBEC3B-like categories. We that used *anz4 MAF files from these filtered mutation catalogs to calculate enrichments with all possible 192 trinucleotide mutational motifs similar to analysis of APOBEC mutagenesis in long persistent subtelomeric ssDNA of yeast hyperexpressing either APOBEC3A or APOBEC3B (see previous section). Mutations in G-strand coordinated clusters were assigned to cytosine deamination in complementary strand

### Knowledge-based statistical valuation of trinucleotide-centered enrichments for RNA edits and DNA mutations to derive signatures

The calculation of the overrepresentation of the mutational or editing motifs was performed using the P-MACD (<u>p</u>atterns of <u>m</u>utagenesis by <u>A</u>POBEC <u>c</u>ytidine <u>d</u>eaminases) pipeline (previously described in ((Chan et al., 2015, 26258849; Roberts et al., 2013, 23852170; Saini et al., 2021, 33444353; (HUDSON *et al*. 2023)) and available at https://github.com/NIEHS/P-MACD)). This P-MACD pipeline calculates the enrichment of a specific trinucleotide motif as compared to random mutagenesis/editing in each Mutation annotation file (MAF) based upon the assumption that a specific motif would occur more frequently among mutated/edited bases than in the surrounding sequence context (context is defined by the +/- 20 bases surrounding the mutated/edited base). The equation to calculate enrichment of a given mutation signature is provided below for the DNA mutational motif tCn to tTn as an example (the mutated nucleotide is capitalized in the motif; n corresponds to any nucleotide).

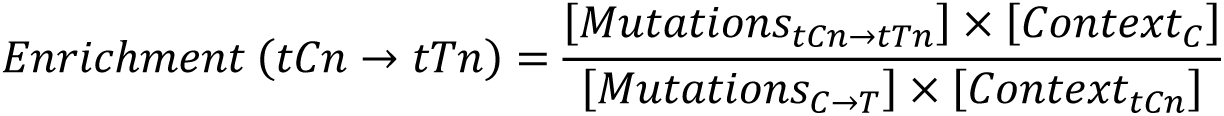

To determine if there is a statistically significant increase in the enrichment for a tested trinucleotide motif, Fisher’s exact test for odds ratios was performed wherein the ratio of the number of mutations within the trinucleotide motif (𝑀𝑢𝑡𝑎𝑡𝑖𝑜𝑛𝑠_𝑡𝐶𝑛→𝑡𝑇𝑛_), and those that do not conform to the trinucleotide motif ((𝑀𝑢𝑡𝑎𝑡𝑖𝑜𝑛𝑠_𝐶→𝑇_) - (𝑀𝑢𝑡𝑎𝑡𝑖𝑜𝑛𝑠_𝑡𝐶𝑛→𝑡𝑇𝑛_)), were compared to the number of bases in the context that were in the trinucleotide motif (𝐶𝑜𝑛𝑡𝑒𝑥𝑡_𝑡𝐶𝑛_) versus those that were not in the motif ((𝐶𝑜𝑛𝑡𝑒𝑥𝑡_𝐶_) - (𝐶𝑜𝑛𝑡𝑒𝑥𝑡_𝑡𝐶𝑛_)).

After MAF files generated for APOBEC-induced ssDNA mutations C to T or RNA editing calls C to U, P-MACD was run for tCa motif common for APOBEC3A and APOBEC3B (CHAN *et al*. 2015). Among P-MACD generated files there were *anz4, which contain all information of the input MAF file plus several additional columns including the column with the +/- 20 bases surrounding the mutated/edited base. This column was essential for calculating enrichments as well as P-values for all possible trinucleotide mutational motifs. When a DNA strand containing nucleotide with a mutagenic lesion is known, enrichment in 192 trinucleotide motifs can be calculated. If a nucleotide change cannot be assigned to one of the two strands, 96 motif enrichments, each totaling enrichments for two reverse complements, are calculated. We calculated enrichments with all possible trinucleotide mutational or editing motifs in ssDNA or in ssRNA, respectively, using the procedure described in (KLIMCZAK *et al*. 2020; HUDSON *et al*. 2023). Since there are only 16 possible C to T or C to U of APOBEC-induced mutations or RNA edits respectively we applied correction of P-values for 16 independent hypotheses using the Benjamini-Hochberg method. For trinucleotide motifs where the enrichment was greater than 1 and the corrected P-value was less than 0.05, odds ratios were compared by Breslow-Day test for homogeneity to evaluate statistical significance of difference between enrichments.

### RNA editing density in secondary structures

The density of RNA editing was calculated as the ratio of C-to-T substitutions within tCn motifs to the total number of potential APOBEC targets, for example the number of tCn motifs in the RNA molecule being analyzed:

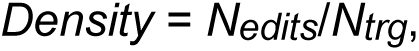

where *N_edits_* represents the number of RNA edits, and *N_trg_* represents the total number of tCn motifs in a target. Calculations for specific positions within a hairpin-loop structure were performed as described in (PONOMAREV *et al*. 2022). Human non-canonical DNA structures, including hairpins, were taken from Non-B DB database (CER *et al*. 2013) For human genome, the calculation of targets was limited to exon regions associated with a transcript of gene where edit(s) were called. For genes producing several transcripts the transcript with the maximum number of edited exons was selected; if multiple transcripts had the same number of edited exons, the transcript with the highest number and the total length of unedited exons was chosen.

## Acknowledgments

This research was supported in part by the NIH, National Institute of Environmental Health Sciences. The contributions of the NIH authors are considered Works of the United States Government. The findings and conclusions presented in this paper are those of the authors and do not necessarily reflect the views of the NIH or the U.S. Department of Health and Human Services. We are grateful to Drs. Paul W. Doetsch, Natalya P. Degtyareva, and Rajula Alleva Elango for advice on the manuscript and for Mr. Adam B. Burkholder for help in data management. D.A.G. is supported by the US National Institutes of Health Intramural Research Program Project Z1AES103266; S.A.R. is supported by R01CA269784 from NCI; M.D.K. is supported by Scientific and Technological Research Council of Turkey (TUBITAK) under the Grant Number 123E476.

## Author Contributions

Conceptualization – D.A.G., Z.W.K., S.A.R.; Data Curation – H.B., Z.W.K., L.P., L.J.K.; Formal Analysis - H.B., L.J.K., Y.C.H., M.D.K.,J.L.L.; Investigation – Z.W.K., M.S.D., M.E.C., T.M.M.; Methodology - D.A.G., Z.W.K., S.A.R.; Resources - L.P., S.A.R.; Software – H.B, L.J.K., M.D.K; Supervision - D.A.G., S.A.R., J.L.L.; Visualization – Z.W.K.; Writing Original Draft - Z.W.K., D.A.G.; Writing – Review & Editing - Z.W.K., D.A.G., S.A.R., H.B.

## Supplementary Information

### Supplementary Figure

**S1 Figure.**
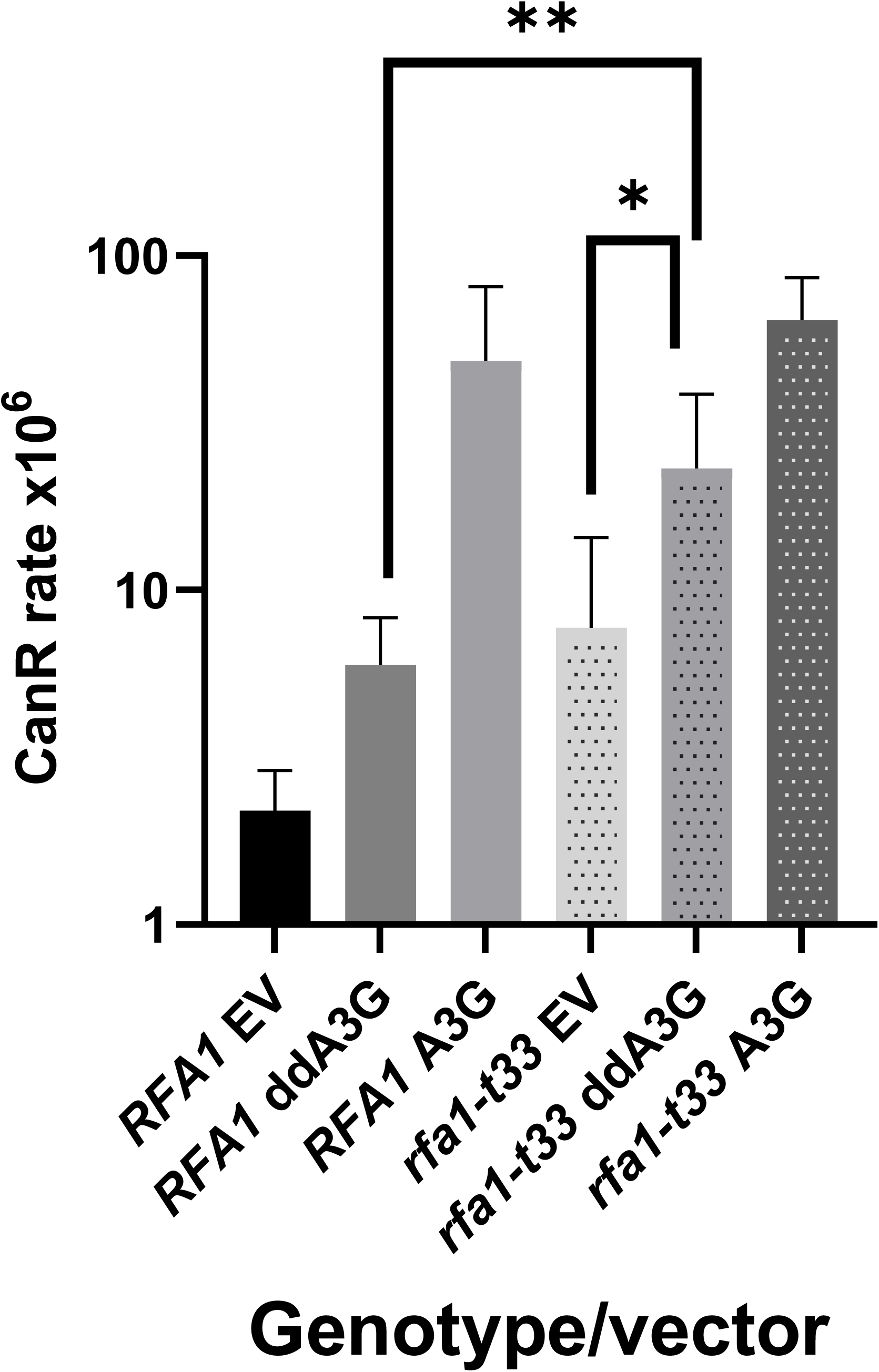
Mutagenesis by single-domain and by full-size double-domain (dd) APOBEC3G in yeast with wild type *RFA1* and hypomorph *rfa1-t33* alleles. CAN1-R mutation rates measured in *RFA1-WT* and *rfa1-t33* strains, carrying either empty vector (EV), single domain APOBEC3G (A3G) or double-domain APOBEC3G (ddA3G) plasmids. Data for EV and single-domain A3G were taken from (DENNEN *et al*. 2024). Shown are median values for mutation rates measured in 6 independent cultures and 95% confidence intervals for the medians. Statistically significant difference with p< 0.05 of the one-tailed Mann– Whitney test showing mutagenic activity of ddA3G in rfa1-t33 background is indicated by brackets. Source data and statistical analyses for all pairwise comparisons can be found in S1 Table.

### Supplementary Tables

**S1 Table**. Comparison of mutagenesis in CAN1 gene caused by single-domain APOBEC3G (A3G) and double domain ddAPOBEC3G in RFA1 and rfa1-t33 yeast strains.

**S2 Table**. Lists of files with DNA mutation and RNA editing calls analyzed in this study Table contains only file lists. Files with DNA mutation and RNA editing lists can be found in: https://doi.org/10.5281/zenodo.18079216

**S3 Table**. Values of motif enrichments used on graphs shown on Figures 1-6

**S4 Table**. DNA motif-centered statistical analyses using 96 trinucleotide-centered mutation motifs

**S5 Table**. Comparison between odds-ratios of different motifs and the same motif in different cohorts

**S6 Table**. DNA motif-centered statistical analyses using 192 trinucleotide-centered mutation motifs

**S7 Table**. Motif-centered statistical analyses using 192 trinucleotide-centered RNA editing motifs Sequences are shown in DNA format (T instead of U) to maintain compatibility with other outputs of mutation signatures

**S8 Table**. Secondary structure preferences of C to U mRNA edits in BT-474 human breast cancer cell line transfected with APOBEC3A or APOBEC3 vector

**S9 Table**. List of accession numbers and publications associated with poliovirus genomes analyzed in this paper

**S10 Table**. Yeast and human cell lines used in this study

**S11 Table**. Detailed documentation of the CLC Genomics Workbench workflow used for calling DNA mutations in bulk yeast cultures expressing strong APOBECs, A3A and A3B

**S12 Table**. Detailed documentation of the CLC Genomics Workbench workflow used for calling DNA mutations in can1 isolated from cultures expressing weaker APOBECs, A1, A3C, ddA3G

